# STEM CELL TECHNOLOGY PROVIDES NOVEL TOOLS TO UNDERSTAND HUMAN VARIATION IN *Plasmodium falciparum* MALARIA

**DOI:** 10.1101/2021.06.30.450498

**Authors:** Alena Pance, Bee Ling, Kioko Mwikali, Manousos Koutsourakis, Chukwuma Agu, Foad Rouhani, Hannes Ponstingl, Ruddy Montandon, Frances Law, Julian C. Rayner

## Abstract

*Plasmodium falciparum* interacts with several human cell types during their complex life cycle, including erythrocytes and hepatocytes. The enuclated nature of erythrocytes makes them inaccessible to genetic tools, which in turn makes studying erythrocyte proteins involved in malaria invasion and development particularly difficult. Here we overcome this limitation using stem cell technology to develop a universal differentiation protocol for *in vitro* derivation of erythrocytes from a variety of stem cell lines of diverse origin. This allows manipulation of erythrocytic genes and examination of their impact on the parasite by flow cytometric detection of parasite haemozoin. Deletion of Basigin, the essential receptor for *P. falciparum*, abrogates invasion, while other less studied proteins such as ATP2B4 have a minor effect. Reprogramming of induced pluripotent stem cells from α-thalassemia primary samples shows reduced infection levels, demonstrating this approach is useful for understanding the effect of natural human polymorphisms on the disease.

## INTRODUCTION

Malaria is an infectious disease caused by several species of *Plasmodium* parasites that are transferred between humans by female Anopheline mosquitoes. The life cycle of the parasite is complex and involves various organs in each host, with a wide range of interactions between parasite and host cell at each step. Nevertheless, the pathology and severe complications of malaria infection in humans result from the blood stages of the infection, during which parasites invade and develop inside erythrocytes. The completion of reference genomes (*1*), the development of genome editing technologies (*2, 3*) and their adaptation to the parasite (*4, 5*) have revolutionised our understanding of the parasite side of the blood cycle. These advances have enabled the identification of many parasite proteins involved in invasion (*6*), as well as the subsequent formation of the parasitophorous vacuole and remodelling of the erythrocyte (*7, 8*), and a much broader understanding of parasite blood-stage biology. On the host side, genome-wide association studies (GWAS) have identified multiple human genetic variants associated with differences in the severity of disease caused particularly by *Plasmodium falciparum*, the most virulent species of parasite affecting humans (*9, 10*). These include many genes implicated in erythrocyte structure and function, such as the membrane protein Band 3, the red blood cell enzyme Glucose-6-Phosphate Dehydrogenase and the Haemoglobins, amongst others (*11*). However, despite population studies providing compelling evidence for protective effects of multiple human protein variants, the molecular mechanisms of these effects are still largely undeciphered. This is mainly due to the difficulties in accessing primary cell samples and the technical challenges of reproducing variants *in vitro* in order to perform tightly controlled cellular studies.

There are two major hurdles for identifying host proteins that interact with malaria parasites and understanding their function. Firstly, erythrocytes are non-proliferative, terminally differentiated cells with a limited life span, making long-term culture of the same cells impossible. Thus, research has relied on clinical samples, with the inherent difficulties of donor availability and variability as well as the impact of storage and transport on sample quality. Perhaps most significantly for mechanistic studies, mature erythrocytes are enucleated and therefore gene editing technologies cannot be applied. These limitations have been partly overcome using siRNA knock-down techniques in Haematopoietic Stem Cells (HSCs) to study the role of Glycophorin A (GYPA) (*12*), Basigin (BSG) (*13*) and to perform a screen for erythrocyte-specific proteins (*14*). However, this approach is curtailed by the restricted availability of HSCs, their limited proliferation capacity and the variable efficiency of the knock-down levels that can be achieved. More recently, an immortalised adult erythroblast line able to proliferate and differentiate *in vitro*, was established by transformation of erythroid progenitor cells with the human papilloma virus HPV16-dervied proteins HPV16-E6/E7 (*15*). One such line (BEL-A) was combined with genome editing technologies to successfully unpick the mechanisms of Basigin involvement in *P. falciparum* invasion (*16*). This approach was also applied to peripheral blood samples and shown to generate cells permissive to *P. falciparum* and *P. vivax* invasion and amenable to genome editing studies (*17*). A disadvantage of this strategy is the viral transformation of the cells with its inherent genetic consequences that might be limiting on the long term, particularly when addressing natural genomic variation.

While these are clearly robust and useful tools, other approaches, such as the use of established Embryonic and induced Pluripotent Stem Cell lines (ESCs and iPSCs) that can be cultured (*18*) and genetically manipulated (*19*) while maintaining their pluripotency (*20*), can provide a versatile alternative with additional advantages. Stem cells have the potential to differentiate into any cell type, which allows examination of the same genomic background as well as any modification or editing on different cell types in all parasite stages. Furthermore, the development of reprogramming techniques that revert terminally differentiated cells to pluripotency (*21*) makes it possible to generate iPSCs from specific individuals. In this way, complex genotypes and rare variants, including non-viable mutations, can be brought into the lab and stored for future studies. Genome editing to change or correct the mutations also become possible, offering a direct confirmation of their physiological role.

In this work we developed a differentiation protocol to drive any stem cell line, ESCs and iPSCs, towards erythropoiesis and produce cells that are competent for *P. falciparum* infection. This approach makes it possible for the first time to study patient-derived cells, while also allowing the potential to incorporate genome editing of specific host genes on multiple different genomic backgrounds. To complete the pipeline we established an *in vitro* assay to accurately quantify invasion of the stem cell-derived erythrocytes by the parasite using an adaptation of flow cytometry based on the refractive properties of haemozoin, a pigment produced by malaria parasites after digestion of haemoglobin. This approach represents a versatile tool to explore a wide variety of host aspects influencing malaria infection and potentially open new avenues for therapeutic intervention.

## METHODS

### Ethics Statement

The use of primary erythrocytes for the culture of Plasmodium falciparum was approved by the NHS Cambridgeshire 4 Research Ethics Committee REC ref. 15/EE/0253 and the Wellcome Sanger Institute Human Materials and Data Management Committee HMDMC 15/076.

The use of human embryonic stem cell lines was approved by the Steering Committee for the UK Stem Cell Bank and for the use of Stem Cell Lines (ref. SCSC11-23) and the Wellcome Sanger Institute Human Materials and Data Management Committee. The Human Embryonic Stem cell lines were obtained from the Centre for Stem Cell Biology, University of Sheffield, Sheffield, UK.

The fibroblast lines from haemoglobinopathy patients were obtained from the NIGMS human genetic cell repository of the Coriell Institute for Medical Research, USA.

### Reprogramming of Induced Pluripotent Stem cell lines

Human IPS lines were derived and verified at the Wellcome Sanger Institute as described (*22-25*). Briefly, 5×10^5^ cells were transduced with Sendai virus carriers of the Yamanaka factors: hOCT4, hSOX2, hKLF4 and hc-MYC overnight at 37^0^C in 5% CO_2_. After a medium change the next day, the cells were cultured for 4 days and from then on maintained in Stem Cell medium: advanced DMEM/F-12 (Gibco, UK) supplemented with 2 mM Glutamax (Gibco), 0.01% β mercapto ethanol (sigma), 4 nM human FGF-basic-147 (Cambridge Bioscience, UK) and 20% KnockOut serum replacement (Gibco, UK), changing medium daily. Ten to 21 days post-transduction, formation of pluripotent colonies was evident, the visible colonies were handpicked and transferred to 12 well plates with MEF feeders. Colonies were expanded into 6 well feeder plates and passaged every 5 to 7 days depending on confluence.

### Cell Culture

All stem cell lines used in this study were cultured on feeder cells (irradiated mouse embryonic fibroblasts MEFS (Global Stem) in the Stem Cell medium described above. The cultures were kept at 37 °C, 5% CO_2_ and medium was changed regularly. Pluripotent cells were passaged using 0.5 mM EDTA (Gibco, UK) and 10µM Rock inhibitor.

### Erythropoietic differentiation

Stem cells were taken off feeder cells with 0.5 mM EDTA (GIBCO) and seeded on gelatin-coated 10 cm plates pre-conditioned with MEF medium over-night and cultured in CDM-PVA supplemented with 12 nM hbFGF (Cell guidance systems, UK) and 10 nM hActivin-A (Source Bioscience, UK). CDM-PVA: 50% IMDM (Invitrogen), 50% advanced DMD-F12 (GIBCO) with 1g/l Poly(vinyl alcohol) PVA (SIGMA), Penicillin/Streptomycin 1x (GIBCO), 1-thioglycerol MTG (SIGMA), Insulin-Transferrin-Selenium 1x (ITS, Life Technologies), Cholesterol 1x (SyntheChol, SIGMA).

As a first step of differentiation, the cells were taken towards the mesoderm germline:

#### Mesoderm 2 Days

CDM-PVA medium supplemented with 5 nM hActivin-A and 2μM SU5402 (SIGMA) <u>Meso/Ery</u>

#### transition 8-12 Days

CDM-PVA medium supplemented with 20ng/ml bFGF, 10nM IL-3 (SIGMA), 10nM BMP4 (R&D Systems), 5μM SB431542 (SIGMA), 5μM CHIR (Axon, The Netherlands), 5μM LY294002 (SIGMA).

During this stage, the detached cells are recovered, washed with PBS and transferred to the erythrocytic differentiation stage, performed in a basic erythrocytic medium (BEM): CellGRO SCGM (CellGenix, Germany) supplemented with ITS, cholesterol, 40ng/ml IGF-1 (Abcam), Penicillin/Streptomycin, 1µM 4-hydroxy 5-methytetrahydrofolate (SIGMA).

#### Ery I 4-5 Days

BEM supplemented with 10ng/ml IL-3, 50ng/ml SCF (Life Technologies), 1µM dexamethasone (SIGMA), 2U/ml EPO (SIGMA), 10ng/ml FLT3 (R&D Systems).

#### Ery II 4-5 Days

BEM supplemented with 50ng/ml SCF, 1µM dexamethasone, 2U/ml EPO

#### Diff Ery minimum 4 Days

BEM supplemented with 2U/ml EPO, 1µM Triiodo-L-Thyronine (T3, SIGMA)

### RNA extraction, qRT-PCR and Microarrays

RNA was extracted using the Isolate II RNA Mini kit (Bioline, UK). 1-3 µg were reverse transcribed with a MuLV reverse transcriptase (Applied Biosystems, UK) using random primers (Bioline, UK). One µl of cDNA was specifically and quantitatively amplified using Biotool 2x SybrGreen qPCR master mix (Stratech, UK) following the cycling parameters established by the manufacturer on a light cycler 480 II (Roche) and using GAPDH as a control for normalisation. The primers used (IDT, Belgium) were:

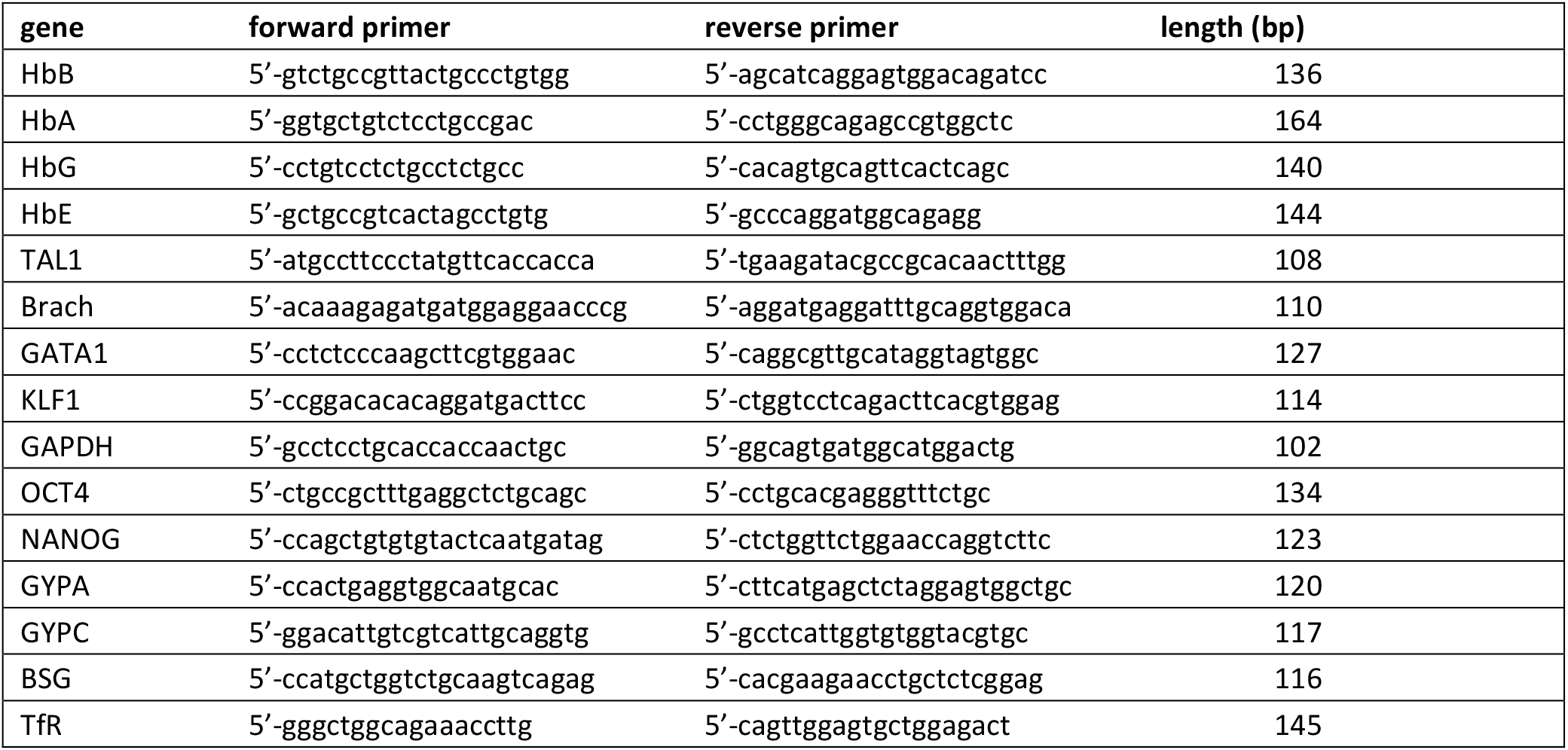

For microarray analyses, RNA was extracted as above, the Illumina TotalPrep RNA amplification kit (Ambion Life technologies) was used to process the samples, and gene expression analysis was assessed on Illumina HumanHT-12v4 chips following the instructions of the manufacturer.

### Parasite culture

Fluorescent *P. falciparum* parasites were cultured in complete RPMI medium (GIBCO) at 2.5% haematocrit with O-RBCs sourced from NHSBT, Cambridge. Cultures were maintained at 37°C in malaria gas (1% O_2_, 3% CO_2_ and 96% N_2_).

### Fluorescent parasites

Parasites were engineered to express a variety of fluorochromes for detection at different wavelengths (*26*). The chosen fluorochromes: tagBFP, Midori-ishi cyan, Kusabira Orange and mCherry were individually inserted into the XhoI / AvrII site of an *attP*-containing vector under regulation by the calmodulin promoter and bearing blasticidin resistance as a selection marker. The NF54 *attB* strain of *Plasmodium falciparum* was transfected with each fluorochrome vector together with an expression vector for Bxb1 integrase and transfectants were selected with blasticidin (2 ugr/ml)(*27*).

### Invasion Assays

Differentiated human stem cells were counted and 1 million cells per assay to be performed were labelled with 1 μM DDAO-HS membrane dye for 2h. After a 30 min wash with medium, the cells were centrifuged (1100 x g for 4’), resuspended in complete parasite culture medium at 75 μl/well and dispensed into a 96 well plate, including a well of cells alone for controls. For controls, primary erythrocytes obtained from the blood bank were labelled in the same way, adding 75 µl at 5% haematocrit per assay.

Asynchronous cultures of fluorescent parasites at a parasitaemia of 1.5 – 2% mature parasites were used to purify schizonts: 10 ml of culture were centrifuged (1100xg for 5’ brake 3), resuspended in 1 ml of medium and loaded onto a 63% Percoll cushion. Centrifugation at 1300xg for 11’ no brake separated the mature parasites at the Percoll interface. These were recovered, washed with parasite medium and resuspended in 3 ml of parasite medium. The parasite suspension, which due to the asynchronous nature of the starting culture contained a relatively broad window of late-stage parasites, from late trophozoites to mature schizonts, was added to the cells in the 96 well plate at 75 μl/well.

One plate per time point was prepared and plates were placed in a gas chamber filled with malaria gas and left in a 37°C incubator for the appropriate length of time.

After the incubation time, the plate was removed, adding 200 μl of PBS/well and spinning at 1100xg 1’. Three μl were taken from the bottom of the wells and smeared on slides to be stained with Giemsa and the supernatant was removed. The pellets were fixed with 100 μl 4% paraformaldehyde for20’, washed and resuspended in PBS to be analysed by flow cytometry.

### Flow Cytometry

Expression of proteins on the membrane of stem cell-derived erythrocytes was measured using specific fluorochrome-tagged antibodies and quantitation by flow cytometry on a LSFORTESSA BD analyser using Flowjo V10.3 (Bcton Dickinson & Co, NJ) and SUMMIT V3.1. Antibodies:

CD71-APC Biolegend #334108

GYPA-PE Southern Biotech #9861-09

BSG-FITC MACS Miltenyi #130-104-489

ATP2B4-FITC LSBio #LS-C446496

A MoFlo flow cytometer (Beckman Coulter, USA) was adapted for detection of laser light depolarisation produced by parasite hemozoin (*28*). The Mo-Flo flow cytometer has a Z-configuration optics platform and is equipped with four solid state lasers (488nm, 561nm, 405nm, 640nm) spatially separated at the stream-in-air flow chamber with 488nm primarily assigned as the first laser. The laser power for 561nm, 405nm, 640nm were all set at 100mW and 488nm was set at 50mW. The laser 488nm, 561nm, 405nm, 640nm was used to excite the Cyan and PE, mcherry, BFP and DDAO respectively. Fluorescence emitted from Cyan, PE, mcherry, BFP and DDAO was collected using a 520/36nm, 580/30nm, 615/20nm, 447/60nm and 671/28nm band pass filter respectively. An optical modification was made on the primary laser detection pod so that the scattered light from 488nm laser light was split into two using a 50/50 beam splitter to measure the normal SSC (vertical) and depolarised SSC (Horizontal) by placing a polarizer (Chroma Technology Corp) with its polarisation axis horizontal to the polarisation plane of the laser light. Both SSC detectors have a 488/10nm band pass filter (Fig. S2). A total of 50000 events was acquired and analysed using Flojo V10.3.

### Genome Editing

Genomic modification to ablate the genes chosen for this study was performed in the RH1 cell line, as described previously (*29*) and shown schematically in supplementary Fig. S8.

### Microarray data analysis

All microarray datasets were put through “neqc” background correction followed by quantile normalization using the limma R package (*30*). Inter-plate variation (batch effects) were adjusted using combat algorithm https://pubmed.ncbi.nlm.nih.gov/16632515/pubmed.ncbi.nlm.nih.gov]. Differential expression analysis was performed to obtain a subset of significant probes (those that change between two or more conditions), FDR adjusted P value of 0.05 was chosen as the cut-off using limma R package. Heatmaps were plotted using Complexheatmap R package https://pubmed.ncbi.nlm.nih.gov/27207943/[pubmed.ncbi.nlm.nih.gov]. The GSE63703 gene expression matrix from GREIN (https://shiny.ilincs.org/grein) was used to identify erythrocyte and erythroid progenitor specific genes.

## RESULTS

### A variety of human stem cells differentiate into erythroid cells *in vitro*

A protocol to differentiate human stem cells into erythrocytes was developed (Fig. S1), in which the adherent pluripotent cells are first taken towards the mesoderm path, as described by Vallier et al (*31*) by exposing the cells to two steps of specific cytokines. As the cells start differentiating, the pluripotency markers decrease while expression of mesoderm markers increases (Fig 1A) followed by the induction of the crucial transcription factors involved in driving the myeloid line of haematopoiesis towards megakaryocyte-erythroid progenitors (MEP). The modulation of gene expression is concomitant with a morphological change of the cells. For our goals, the final stage of this phase was extended to 8 – 12 days during which the cells detach spontaneously from the plate.

**Figure 1A.**
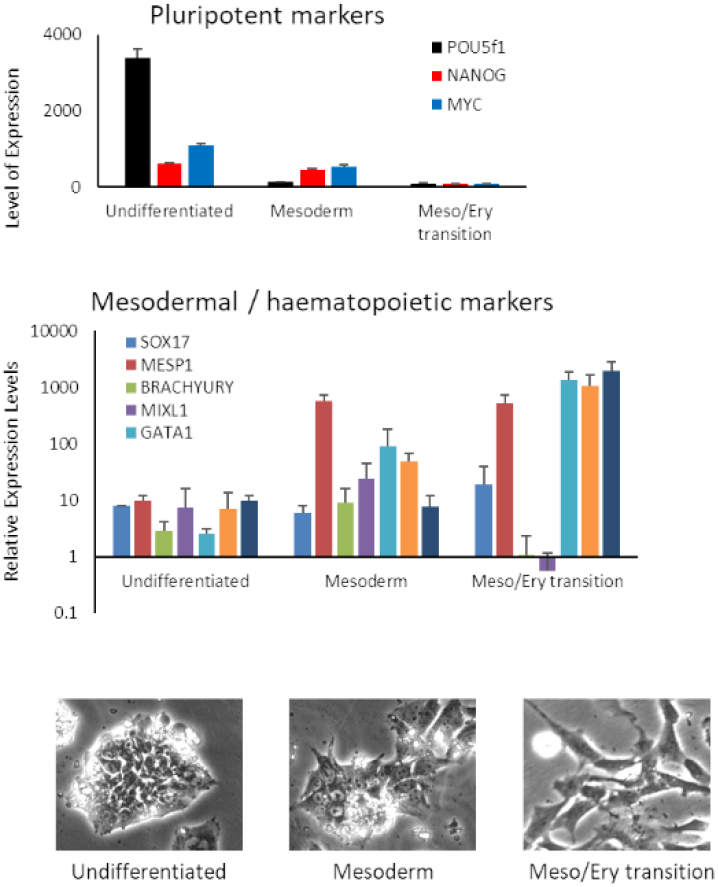
Differentiation of stem cells towards erythropoiesis. **A:** Mesoderm stages of differentiation. Expression of pluripotency (POU5f2 (Oct 4), Nanog and c-myc), mesoderm (Sox17, MESP1 and Brach) and haematopoiesis (Mixl1, GATA1, Tal1, KLF1) markers, as measured by qRT-PCR and morphological change of the differentiating cells is documented by light microscopy (1000x).

The suspension of cells detaching from the plate is then taken through the erythropoietic phase of the protocol during which the cells are exposed to Erythropoietin (EPO) and IL-3 to drive differentiation into erythroblasts. The cells are kept in dexamethasone in order to halt the process at the erythroblast stage and homogenise the culture. Next, the erythroblasts are expanded and differentiation is completed by removing dexamethasone (Fig. 1B). At the end of the differentiation process, analysis of the characteristics of the cells show surface expression of Glycophorin A (GYPA), a major marker of erythrocytes and expression of adult haemoglobins A and B (Fig. 1B). The cells also express the transferrin receptor (CD71) and foetal haemoglobins Gamma and Epsilon and the great majority of them (70 to 80%) stain positively with a DNA dye (Hoechst33342), indicating that they still have some form of nucleus or nucleic acid content (Fig 1B). These features indicate that the cells generated correspond to immature erythroid cells earlier in the differentiation pathway to mature erythrocytes. The full transcriptome changes as the cells differentiate were analysed using microarrays, showing increasing expression of erythrocytic genes including membrane proteins, components of the haeme cycle and the haemoglobins (Fig. 1B). The microarrays also confirmed that extension of the mesoderm/erythroid transition step of the mesoderm stage improves erythrocytic differentiation and that a duration of 8 or 12 days leads to similar gene expression patterns.

**Figure 1B.**
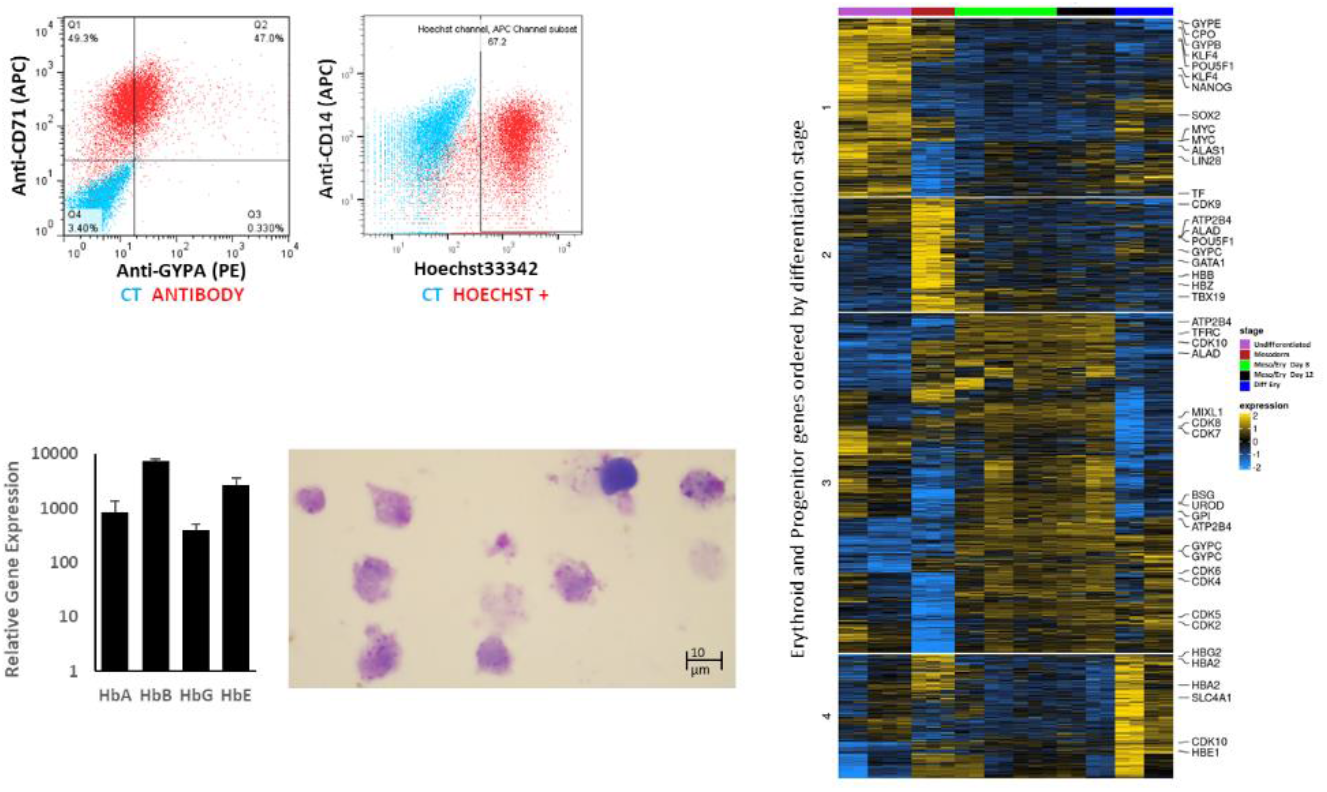
Final erythropoietic stage of differentiation. Expression of the erythrocyte markers: Glicophorin A (GYPA) and transferring receptor (CD71), and DNA labelling with Hoechst33342 were assessed by flow cytometry; expression of the haemoglobins was quantified by qRT-PCR. Morphology of the cells is shown by Giemsa staining and light microscopy (1000x). An overall examination of erythrocytic and erythroid progenitor gene expression throughout the differentiation protocol was assessed by microarray analysis.

To establish the versatility of the differentiation protocol, a variety of cell lines of different origins were tested. This included human embryonic stem cell (hESC) lines (Shef3 and Shef6), as well as Induced Pluripotent Stem Cell (hIPSC) lines derived from fibroblasts (RH1, SF2 and K4) and from blood (CD3, CD5 and GB1, GB4). The efficacy of the process was assessed according to the expression of surface markers GYPA and CD71 by flow cytometry (Fig. 2A) and the haemoglobin genes by qRT-PCR (Fig. 2B). Figure 2 shows that though there are variations between the lines tested, they all differentiate in a similar fashion. The overall expression of erythroid genes revealed by microarray analysis confirmed a similar response by all cell lines tested to the differentiation protocol (Fig. 2C).

**Figure 2.**
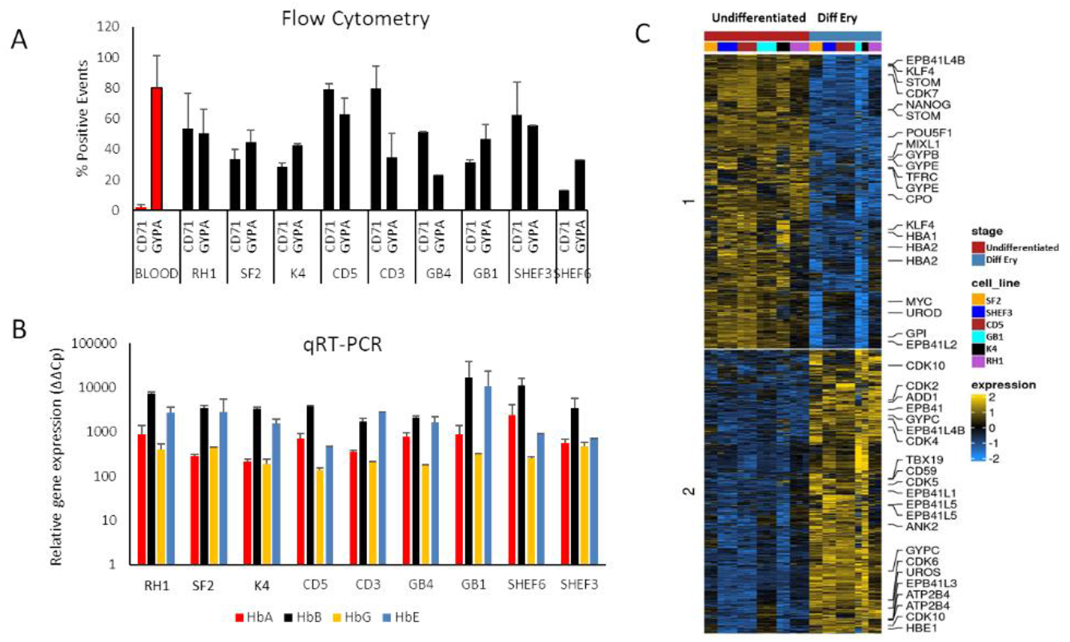
Characterisation of a variety of stem cell lines differentiated to erythrocytes in vitro. **A:** erythrocytic surface markers Glycophorin A (GYPA) and the transferrin receptor (CD71) by flow cytometry. **B:** expression of the haemoglobins (adult A and B; foetal G and E) assessed by qRT-PCR. **C:** comparison of gene expression by microarray analysis at Undifferentiated and final erythropoietic differentiation stage.

### Stem cell-derived erythroid cells support invasion by *Plasmodium falciparum*

The capacity of the different cell lines to support *P. falciparum* infection was assessed with a purposely developed assay. The *in vitro* differentiated cells are fluorescently labelled to distinguish them from any primary erythrocytes co-purified with the parasites, and cultured with percolled parasites from fluorochrome-expressing *P. falciparum* lines (Fig. S1). The use of purified fluorescent late-stage parasites ensures prompt release of merozoites for invasion and avoids the traditional labelling with DNA dyes that is incompatible with the presence of nucleic acids in the in vitro-derived erythrocytes. To detect invasion, we used a flow cytometry approach specifically adapted to detect haemozoin (Fig S3), the metabolic product of the parasite’s digestion of haemoglobin (*28*). The strategy for the quantification is shown in Figure S4: briefly, after eliminating debris and doublets, the infected differentiated erythrocyte population is selected based on the fluorescence of both cells and parasites, and then examined for depolarisation using uninfected cells to establish the gates. Since light depolarisation is an exclusive property of haemozoin produced by the parasite, the events detected correspond to parasites that successfully invaded labelled *in vitro*-differentiated cells. Because the quantification of depolarising events is applied to the selected population of labelled cells, the proportion reflects the parasitaemia of the culture. To determine the optimal combination of fluorescent labels for the assay, parasites expressing a range of fluorochromes were generated (*26*) and different cell dyes were tested. Invasion of cells labelled with the green cell dye DFFA with mCherry-parasites (Fig. S4) was equally effective for quantification of parasitaemia as labelling cells with the far-red dye DDAO and infecting them with Midori-ishi cyan- or tagBFP-parasites (Fig. S5 and S6). The combination of DDAO-stained cells and tagBFP parasites was chosen for all following experiments because these two labels are very strong with the best separation of their excitation/emission spectra.

Invasion is first quantified after 18 hours to allow enough time for haemozoin accumulation and fluorescent protein expression. As haemozoin synthesis increases throughout the blood cycle, a second time point towards the end of the erythrocytic cycle at 42 hours is also measured to assess parasite growth. Examination of Giemsa-stained slides under the microscope confirmed successful invasion (Fig 3A). All cell lines differentiated were effectively invaded by *P. falciparum* reaching a parasitaemia between 5 and 8% (Fig. 3B), similar to the levels observed in control assays with labelled blood (4.55%) (Fig. S7A). At 42 hours parasitaemia rose in all cell lines to 7–10%, reflecting the growth of small parasites that went undetected at the 18 hour timepoint (Fig. 3B). The blood controls showed a significantly higher increase in parasitaemia at the 42-hour time point (*P*<.001) with an average of 20% (Fig. S7A), likely due to the much higher cell-to-parasite ratio in these control wells. The purified parasites contain a range of late stages from late trophozoites to mature schizonts, meaning that some parasites will mature and rupture within a 10-12 hour window after the set-up of the assay (see Methods section). The 2.5% haematocrit of the blood assays represents an approximate ratio of 30:1 erythrocytes to parasites offering ample availability for continuous invasion throughout the assay. By contrast, because the number of *in vitro*-derived erythrocytes is limited, these assays have a ratio close to 1:1, restricting invasion to the initial merozoites released in the assays.

**Figure 3:**
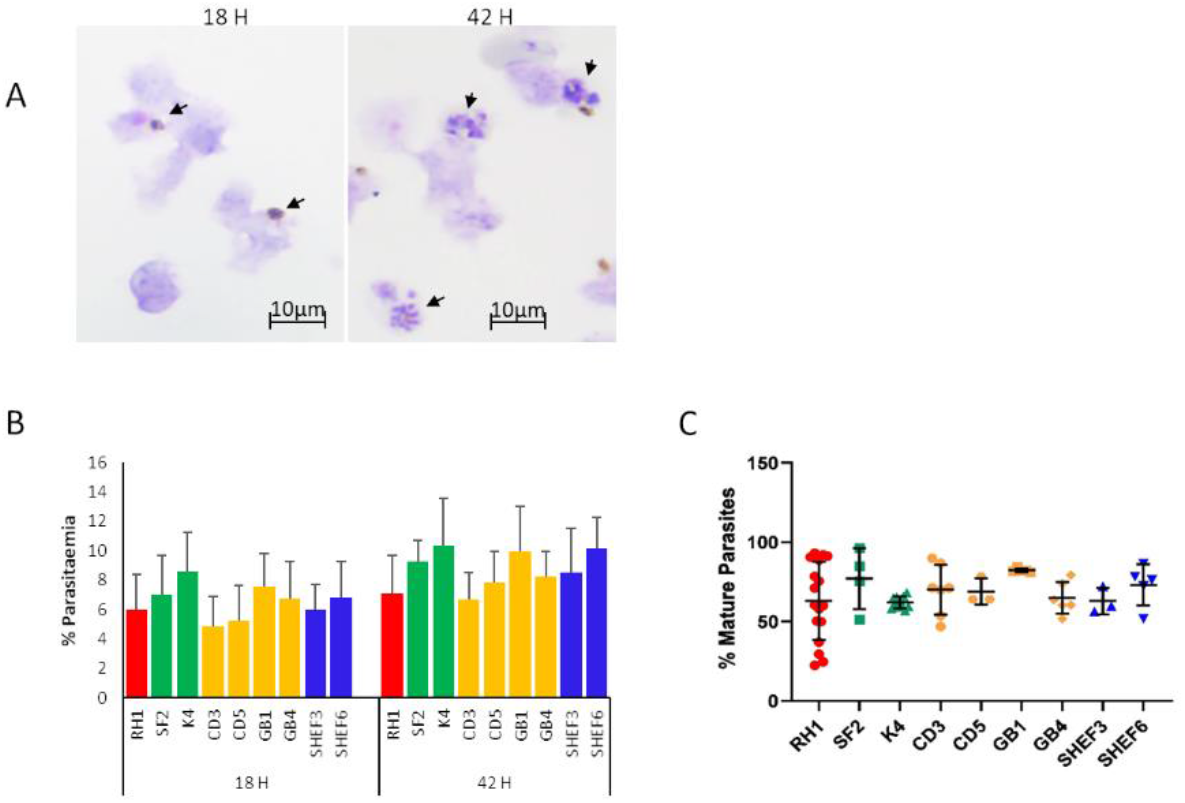
Invasion of in vitro-generated erythrocytes with P. falciparum. **A:** example of Giemsa staining of invasion assays on slide smears, parasite are indicated with arrows. **B:** quantification of invasion by flow cytometry of a variety of differentiated cell lines at two time points of culture, 18h (invasion) and 42 hours (development). **C:** quantification of mature parasites percentage at 42 hours.

The concomitant increase in haemozoin intensity with life cycle progression is a useful tool to measure parasite growth (*28, 32*). Indeed, synchronised parasite cultures show a clear shift in haemozoin signal intensity as they progress from rings to schizonts (Fig. S7B), and this is replicated in our bespoke invasion assays (Fig. S7C), albeit a wider 42-hour peak resulting from the characteristics of the assays as explained above. The gate established by the shift in haemozoin intensity between the ring and schizont stages (M1 in Fig. S7B), was used to quantify the percentage of mature parasites in the cultures (Fig. S7C), which showed an average of 64% (Fig. S7D). Applying the same gate to the assays with the *in vitro*-generated erythrocytes, we observed similar levels of mature parasites between 60 and 80% (Fig. 3C).

### Genome editing of human stem cells reveals a role for specific genes in malaria invasion

The potential to derive edited erythrocytes with this approach was tested with the control line RH1 using CRISPR/Cas9 technology (Fig. S8). Two target genes were chosen, Basigin (*BSG*, CD147) which is known to be a universal receptor for *P. falciparum* invasion (*13*) as a proof of principle, and *ATP2B4* (*PMCA4*) since natural variation in this gene has been correlated to resistance to severe malaria (*33*). The edited clones were genotyped (Fig. S9A) and verified by sequencing (Fig. S9B) confirming small deletions in the critical exon that generate a stop codon downstream. The lack of protein expression was confirmed with specific FITC-labelled antibodies for flow cytometry (Fig 4A).

**Figure 4:**
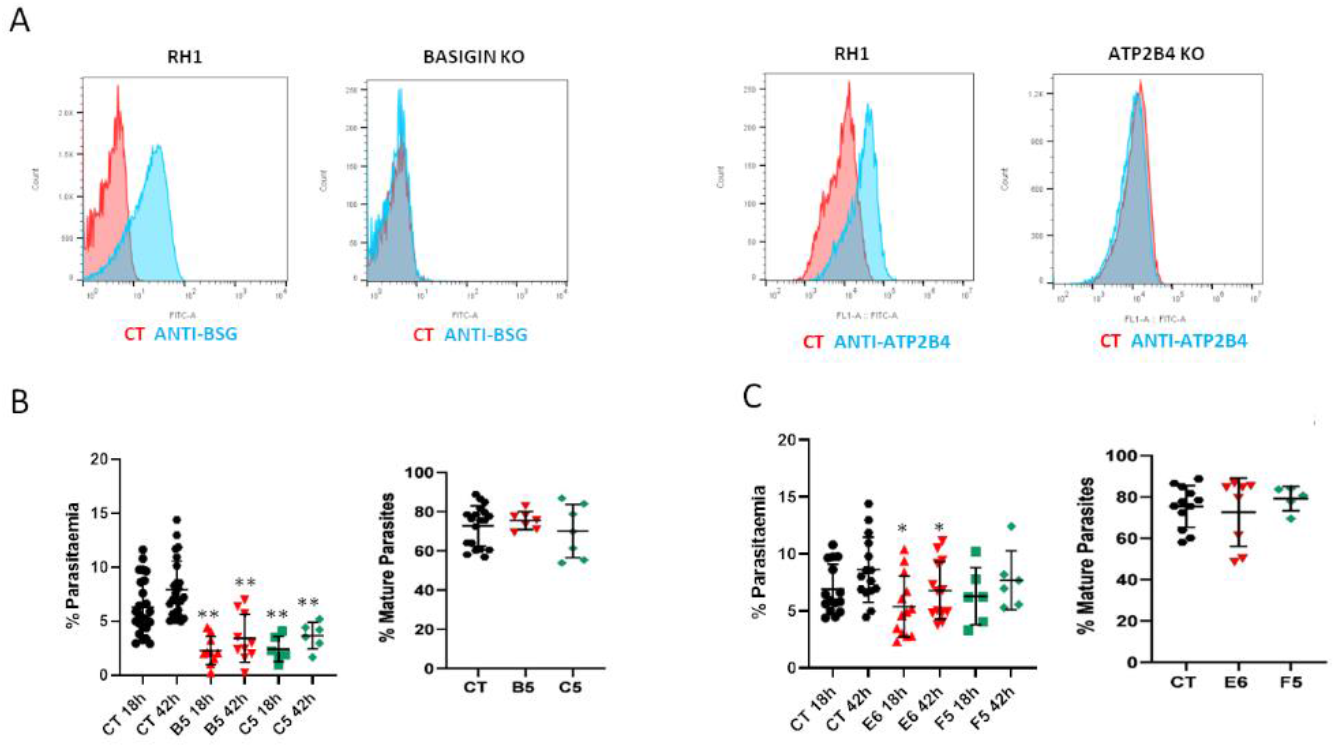
Gene editing in stem cells to study specific genes: Basigin and ATP2B4. Each gene was deleted using CRISPR/Cas9 and clones were isolated. **A:** expression of the protein corresponding to the deleted gene was measured on the cell surface with specific antibodies by flow cytometry. **B:** invasion of Basigin KO clones B5 and C5 ** P<.001 and percentage of mature parasites at 42 hours. **C:** invasion of ATP2B4 KO clones E6 and F5 * P<.05 and percentage of mature parasites at 42 hours.

When challenged with the parasites, neither of the independent Basigin-null cell lines (B5 and C5) supported invasion (Fig 4B), consistent with its role as an essential receptor for *P. falciparum* (*13*). The proportion of mature parasites after 42 hours of culture was similar to the control cells (Fig. 4B), indicating that the few parasites that invaded the modified cells could achieve some growth. Deletion of ATP2B4 (Fig. 4A) showed a tendency to lower invasion levels, marginally significant for only one clone (Fig 4C). The proportion of mature parasites in the ATP2B4 KO cultures at the 42-hour time point was similar to that observed in the control lines, indicating that disruption of this gene does not have a major effect on the development of the parasite.

### Reprogramming IPS lines from haemoglobinopathy patients shows the versatility of this system

Fibroblast lines isolated from patients with α-thalassemia (HbBart) haemoglobinopathies were sourced from the Coriell repository and reprogrammed to generate IPS lines (Coriell Cat# GM10796, RRID:CVCL_N352 called Euml (EM); Coriell Cat# GM03433, RRID:CVCL_N008, called Fijo (FJ)). The IPS lines obtained were differentiated in parallel with our reference cell lines and exposed to fluorescent parasites in our invasion assays. As shown in Fig 5, both thalassaemic cell lines showed significantly decreased efficiency of invasion. Furthermore, the proportion of later stages of parasites in these cultures at the 42-hour time point was also reduced compared to the control cell lines (Fig. 5).

**Figure 5:**
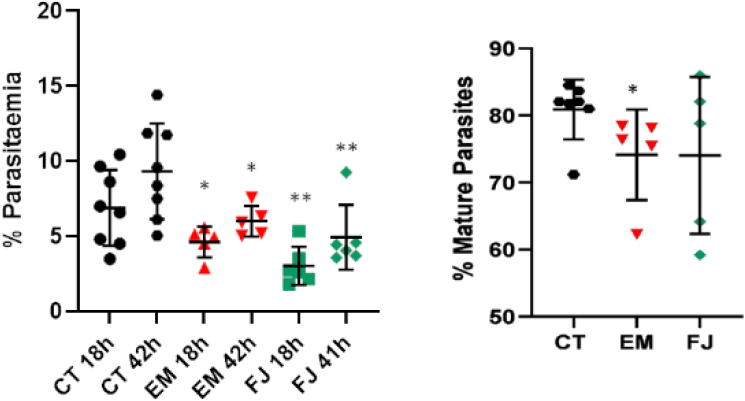
Reprogramming of IPS cells from patients with α- thalassemia haemoglobinopathy. Fibroblasts from α- thalassemia major samples were reprogrammed to IPS cell lines that were differentiated and exposed to P. falciparum to assess invasion and percentage of mature parasites at 42 hours * P<.05; ** P<.005.

## DISCUSSION

In this work we present a protocol that effectively differentiates a variety of human stem cell lines towards erythropoiesis and generates cells that are susceptible to infection with *Plasmodium falciparum* parasites. A total of 9 stem cell lines of diverse origin were studied, showing that though the differentiation efficiency does vary slightly between lines, they all generate erythroid cells as demonstrated by upregulation of erythrocytic genes and expression of erythrocytic proteins. Despite enucleation being notoriously difficult to achieve *in vitro* (*34, 35*) we observed enucleation levels of 20-30%. In fact, the cells with positive nucleic acid staining also include cells with nuclear fractionation and some in which extrusion of the nucleus is incomplete, as can be seen in Figure 1B. Importantly, stem cells differentiated with this protocol are capable of supporting invasion by *P. falciparum* without the need to sort or purify the differentiated cells.

During the blood cycle, haemoglobin is the main source of amino acids for the parasite’s metabolic needs. Degradation of haemoglobin releases free haem, which represents a major toxic insult to the parasite. In a detoxification mechanism the oxidised iron group is compacted into an insoluble crystalline form: β-hematin or haemozoin and stored in the food vacuole (*36*), becoming a distinct feature of intra-erythrocytic Plasmodium parasites (*32*). Intracellular haemozoin formation is reliably detected 12 to 18 hours post-infection and both crystal size and number increase with progression of the blood cycle (*37, 38*). The flow cytometry strategy to detect depolarisation of light caused by hemozoin therefore ensures that the events detected correspond to parasites that have successfully invaded differentiated cells, survived and started growing. However, because haemozoin only becomes detectable at the late ring stage, smaller rings can be missed at the first time point of 18 hours. The use of schizonts for the assays ensures quick invasion after the co-culture is set up, but complete synchrony is difficult to achieve as there will always be a contribution of less mature parasites in the schizont preparations, particularly from routine asynchronous cultures. Therefore, the increase in the number of haemozoin-positive events at 42 hours is likely due to the maturation of late-invading parasites that become detectable at the later time.

The successful manipulation of genes implicated in malaria infection was demonstrated by deletion of *Basigin*, which resulted in a dramatic decrease of infection as expected given the known role of this protein in invasion (*13*). Natural variants in ATP2B4 have been associated with resistance to malaria in various studies (*10, 39*), but the mechanism of protection is not known. A number of variant SNPs have been identified in this gene, mostly in Linkage Disequilibrium (LD), and though it is not clear whether all these SNPs play a role in protection against severe malaria, one of them was shown to disrupt a GATA-1 site in the promoter of the gene (*40*). As a consequence, the expression levels of the protein are markedly reduced giving rise to changes in erythrocyte parameters such as mean corpuscular haemoglobin concentration (MCHC) and size. ATP2B4 is the main membrane Calcium ATPase of erythrocytes that removes calcium from the cytosol to maintain the low levels necessary for calcium-dependent signalling to occur (*41*). A role of calcium in the invasion process of *P. falciparum* has been suggested (*42, 43*) and it is also possible that impairment of calcium homeostasis will affect survival and development of the parasite in the erythrocyte (*44*). Furthermore, reduced levels of ATP2B4 lead to a reduced volume and MCHC (*40, 45*), and these parameters are also predicted to impact the development of the parasite. However, a knock-out of ATP2B4 in our system did not show a major effect on *P. falciparum* invasion or on growth, though a tendency towards a reduction in both parameters was observed. A compensatory effect of PMCA1, which represents 20% of erythrocytic Calcium ATPases, could explain the minimal effect of deleting ATP2B4. It is also possible that the natural impact of ATP2B4 variation on *P. falciparum* growth is relatively subtle and cannot be detected in only one erythrocytic cycle as measured in our assays, but could make the tendency observed important over the time of an infection. Alternatively, as *ATP2B4* is widely expressed throughout the body, variations in the gene might affect other aspects of the disease, such as the interaction of infected erythrocytes with endothelial cells or with the brain, as has also been suggested (*33*).

We further demonstrate the adaptability of this strategy by reprogramming iPS cells from haemoglobinopathy patients. For this we chose one of the traits that is known to confer protection against malaria: thalassemia. Alpha-thalassemia results from a variety of large deletions affecting one or more of the duplicated alpha globin genes and the severity of the disease depends on how many of the four genes are affected. Loss of all 4 α-globin genes, known as α-thalassemia major, can occur in the common South East Asian deletion, leading to the lethal HbBarts hydrops foetalis. Alpha-thalassemia major was chosen for these studies because of the extreme phenotype and also because primary erythrocytes with this genotype are unavailable to perform laboratory assays, thus highlighting the advantages of stem cell technology. Both of our derived reprogrammed cell lines are null for alpha globin, presenting –SEA/–SEA (GN03433) (*46*) and – SEA/–Fil (GM10796) (*47*) genotypes. When differentiated, both cell lines showed a significantly reduced ability to support *P. falciparum* infection, consistent with reported effects of haemoglobinopathies on malaria (*48, 49*). Though several mechanisms have been proposed, it is still unclear how haemoglobin deficiencies impact the parasite and their study is complicated by the variety of genetic changes underlying these traits as well as the difficulty in obtaining samples of primary erythrocytes. It is known that the imbalance in the synthesis of globin chains in alpha and beta thalassemias result in impairment of the assembly of haemoglobin tetramers. This leads to the formation of haemoglobin precipitates (Heinz bodies), which together with the increased hydration occurring in α-thalassemias impair erythrocyte deformability (*50*). It was shown that erythrocyte deformability is lower in samples of α-thalassemia traits in which 2 alpha globin genes are inactivated and the decrease is much stronger in Haemoglobin H disease in which 3 alpha globin genes are missing. Furthermore, this decrease in deformability was directly corelated to decreasing *P. falciparum* invasion (*51*). It is reasonable to predict an even greater deformability defect in the total absence of haemoglobin alpha of the cell lines used here, which is consistent with the dramatic decrease in invasion we observed. Additionally, it was shown that *P. falciparum* parasites produce significantly lower numbers of merozoites in alpha and beta thalassemia trait cells, which correlates with the MCHC and mean corpuscular volume (MCV) of these cell types (*52*).

The strategy presented here is based on a universal differentiation protocol that allows the *in vitro* derivation of erythrocytes from a variety of stem cells lines amenable to the study of host factors involved in malaria infection. Stem cells of any origin can be used, including existing resources of ESC and iPSC lines and also offers the possibility to generate and study novel patient-derived iPS lines. The latter is a novel application of stem cell technology for the study of malaria that presents the only option to study complex genetic traits as well as multiple mutations in their full genomic context. Genome editing can also be applied to introduce or correct specific changes to identify human factors involved in the disease and understand the mechanisms of the impact of genomic variation. The cell lines generated have unlimited proliferation potential and can be stored, thus preserving the genetic characteristics. They also have the potential to be differentiated into a variety of cell types allowing examination of the same genotypes in different stages of the disease. This approach is a powerful tool for the understanding of this disease, circumventing limitations such as availability and access to primary cells with certain traits and complex polymorphisms. A universal differentiation protocol such as we present here greatly increases the versatility and power of this stem cell-based system for a wide range of applications for future malaria research.

## SUPPLEMENTARY FIGURES

**Figure S1.**
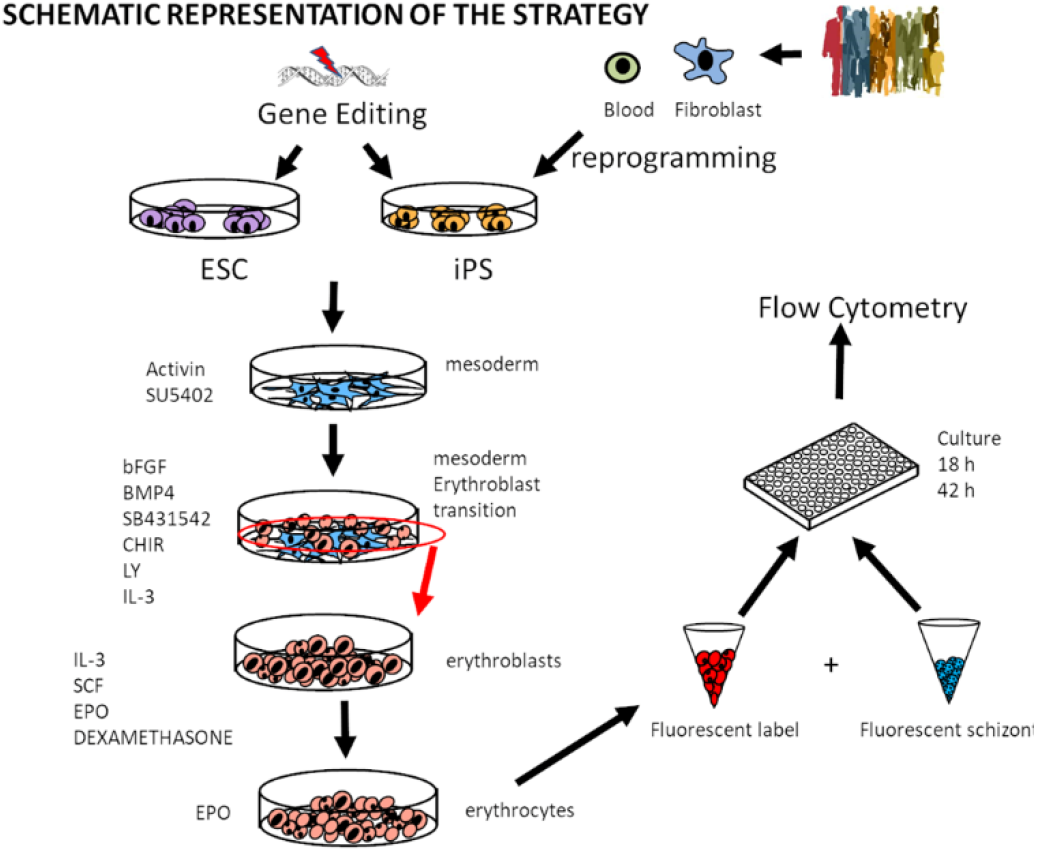
Protocol for Stem cell differentiation. Stem cells can be reprogrammed or edited and differentiated into erythrocytes and then challenged with fluorescent parasites to assess invasion and growth.

**Figure S2.**
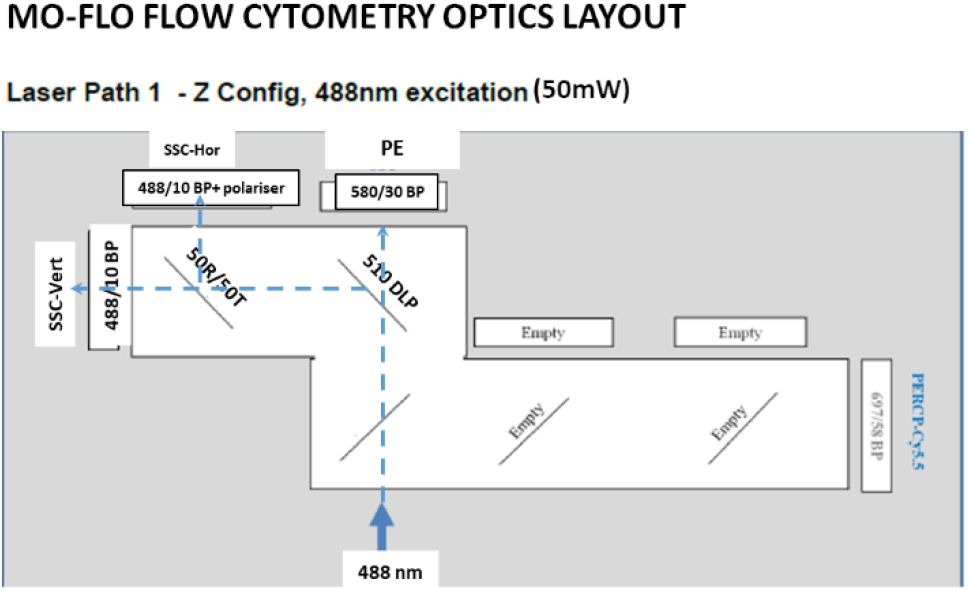
Flow cytometer setting. Setting of the he MO-FLO cytometer in order to detect light depolarisation by haemozoin.

**Figure S3.**
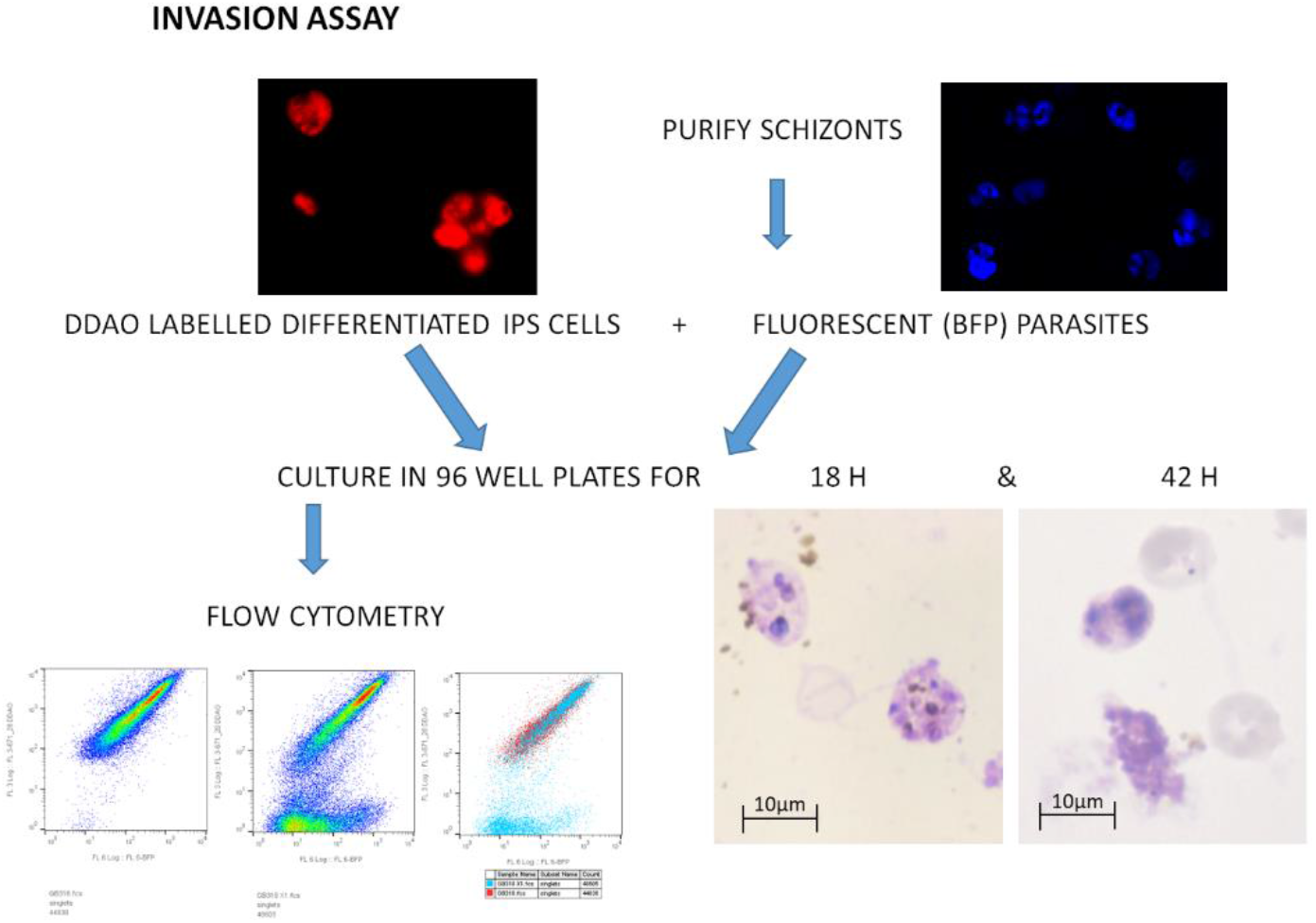
Invasion assay. Labelled differentiated cells are incubated with late stage fluorescent parasites for 18 h and 42 h. The cultures are fixed with PFA (4%) and assessed by flow cytometry.

**Figure S4.**
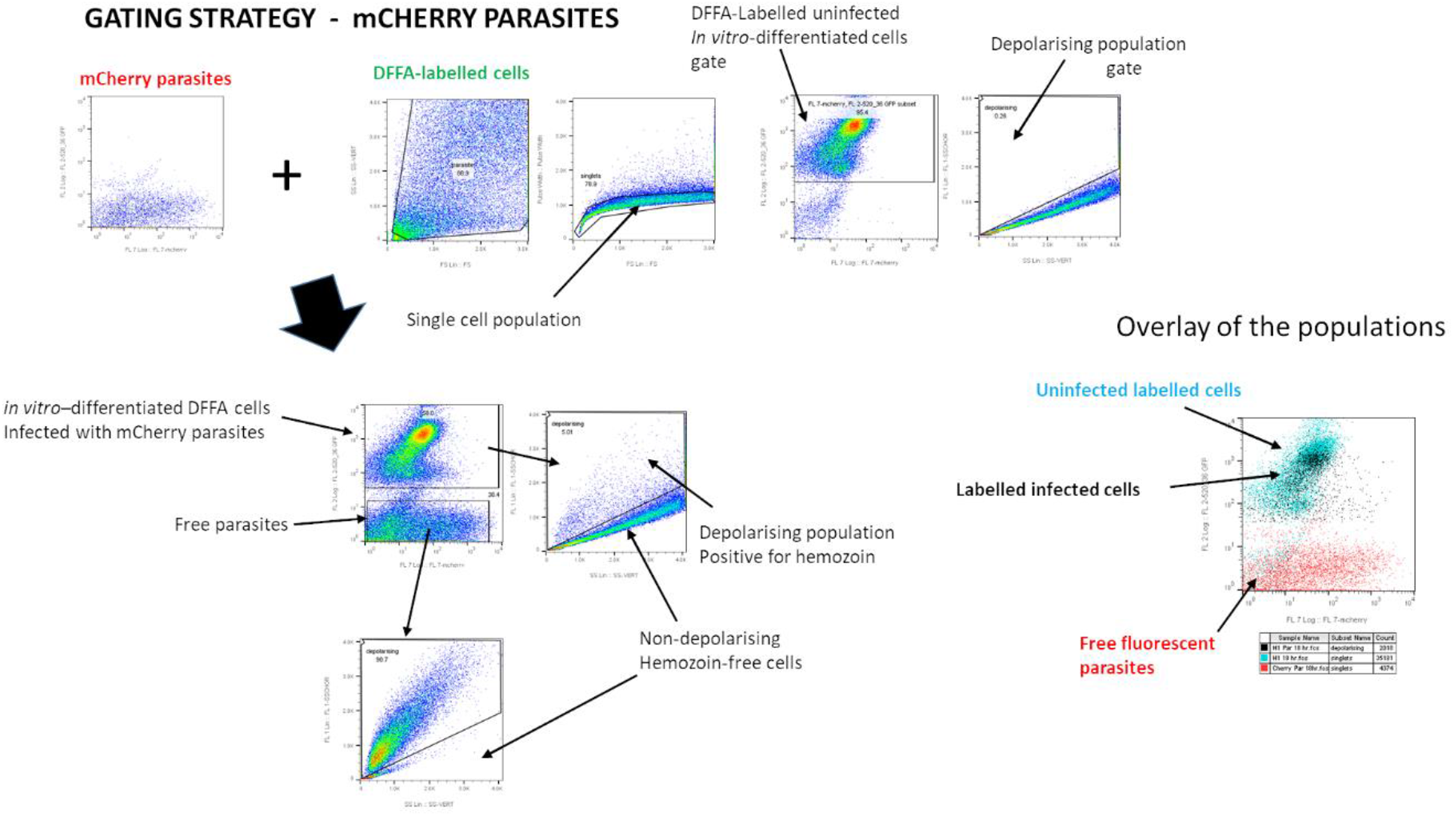
Flow cytometry gating strategy. The population of labelled cells is separated from debris and doublets are eliminated. The fluorescent single cell population is assessed for depolarisation and the gate is established with uninfected cells. The same process is followed for the co-cultures which reveal the haemozoin positive events that are confirmed as infected cells by superimposition of the populations.

**Figure S5.**
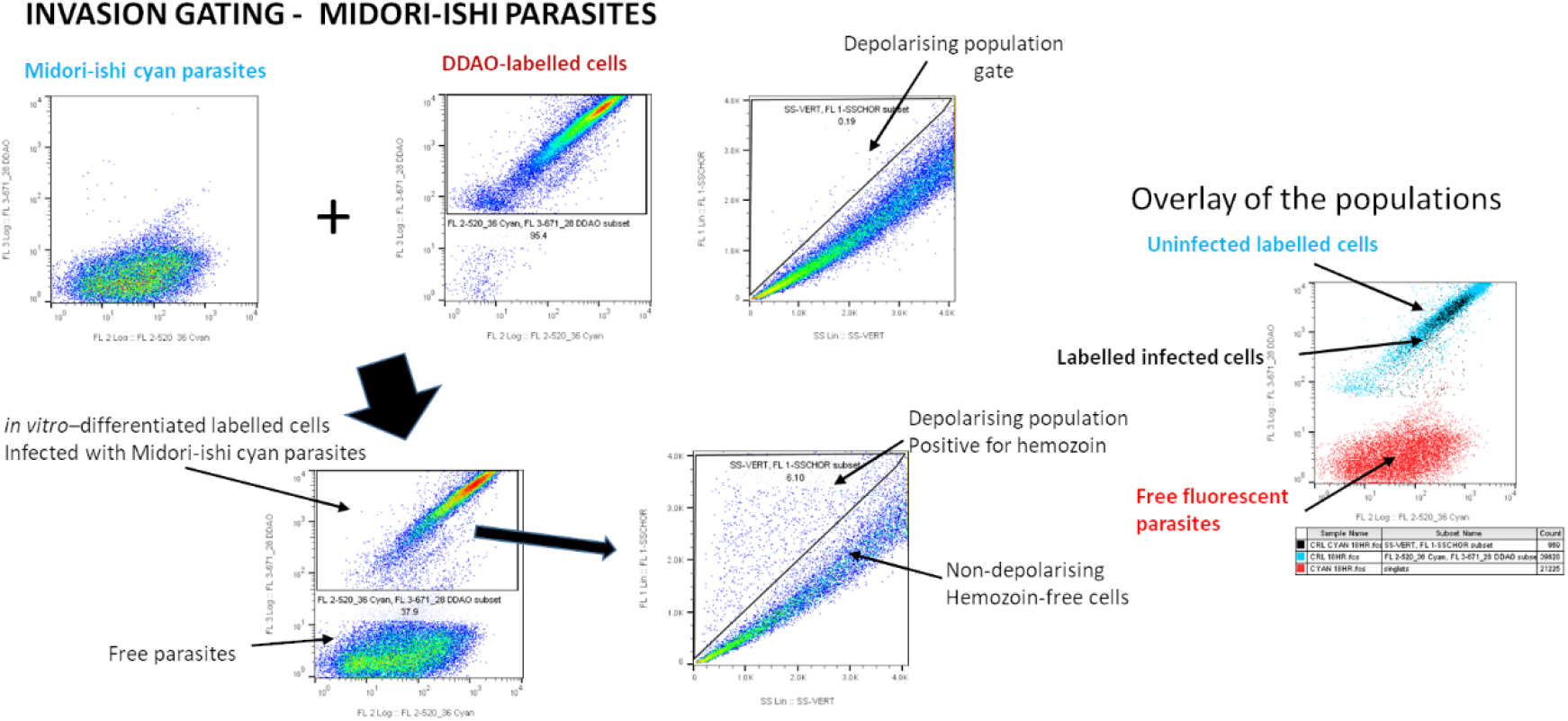
Invasion with Midori-ishi cyan fluorescent parasites.

**Figure S6.**
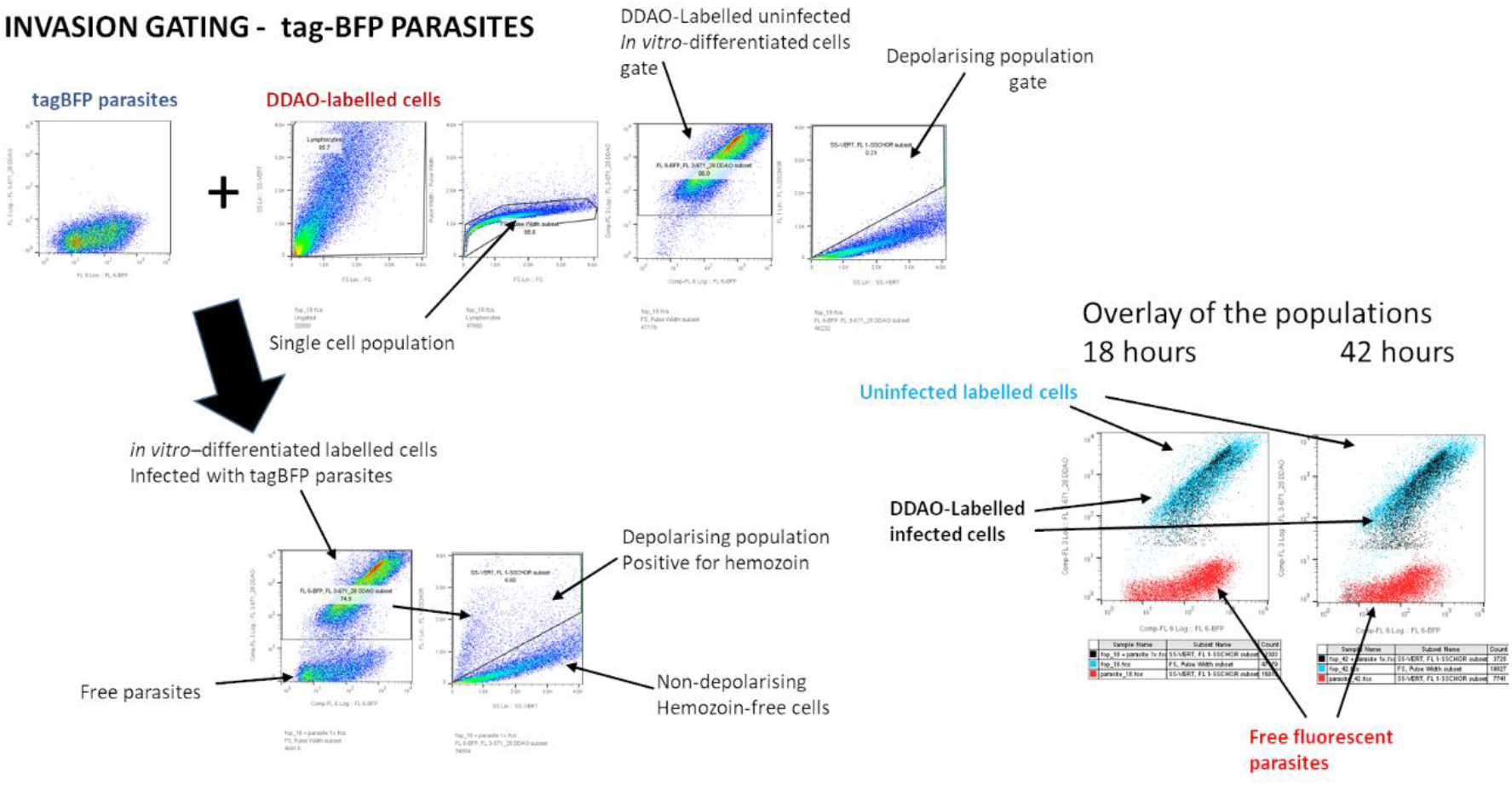
Gating of invasion assays with tag-BPF fluorescent parasites.

**Figure S7.**
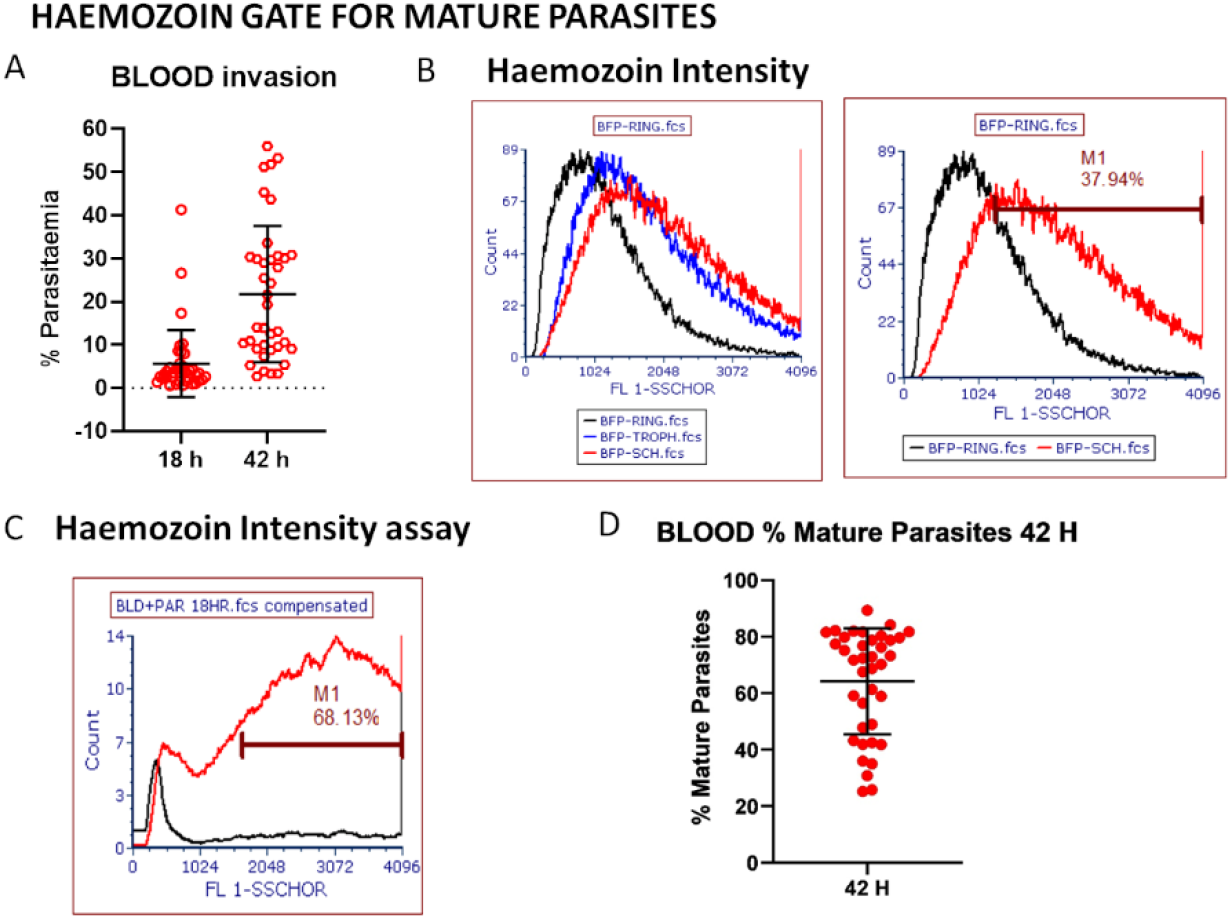
Invasion of blood controls. Every assay had control wells with labelled blood that were analysed in the same way as in vitro derived erythrocytes (A). Synchronized parasites were used to assess the shift in haemozoin intensity (B), which was also measured in the regular assays (C and D).

**Figure S8.**
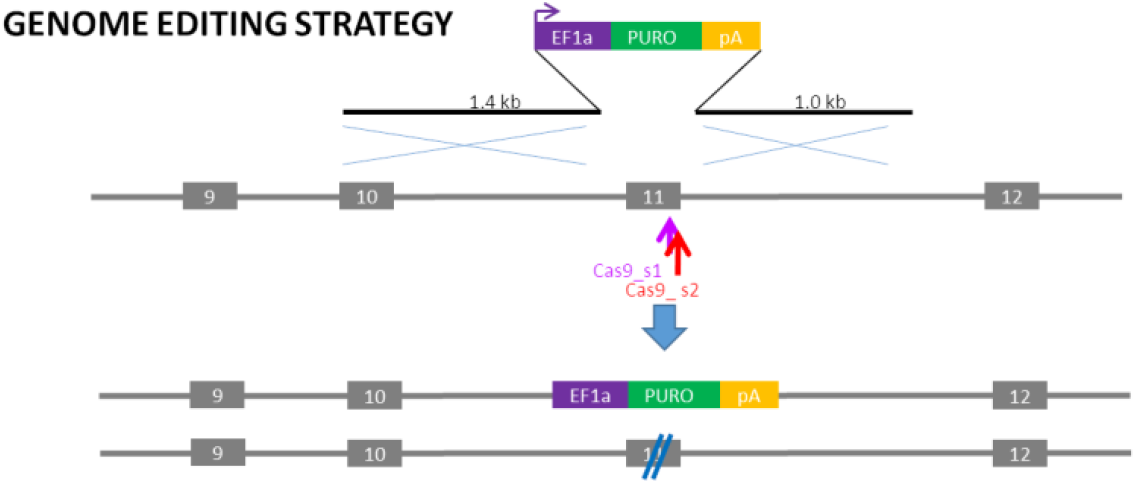
CIRSPR/Cas9 editing of iPS cells. The critical exon was identified as present in all isoforms and creating a stop codon or frame shift when deleted. Vectors were constructed to delete this exon inserting a selection cassette in one allele and disrupting the other.

**Figure S9.**
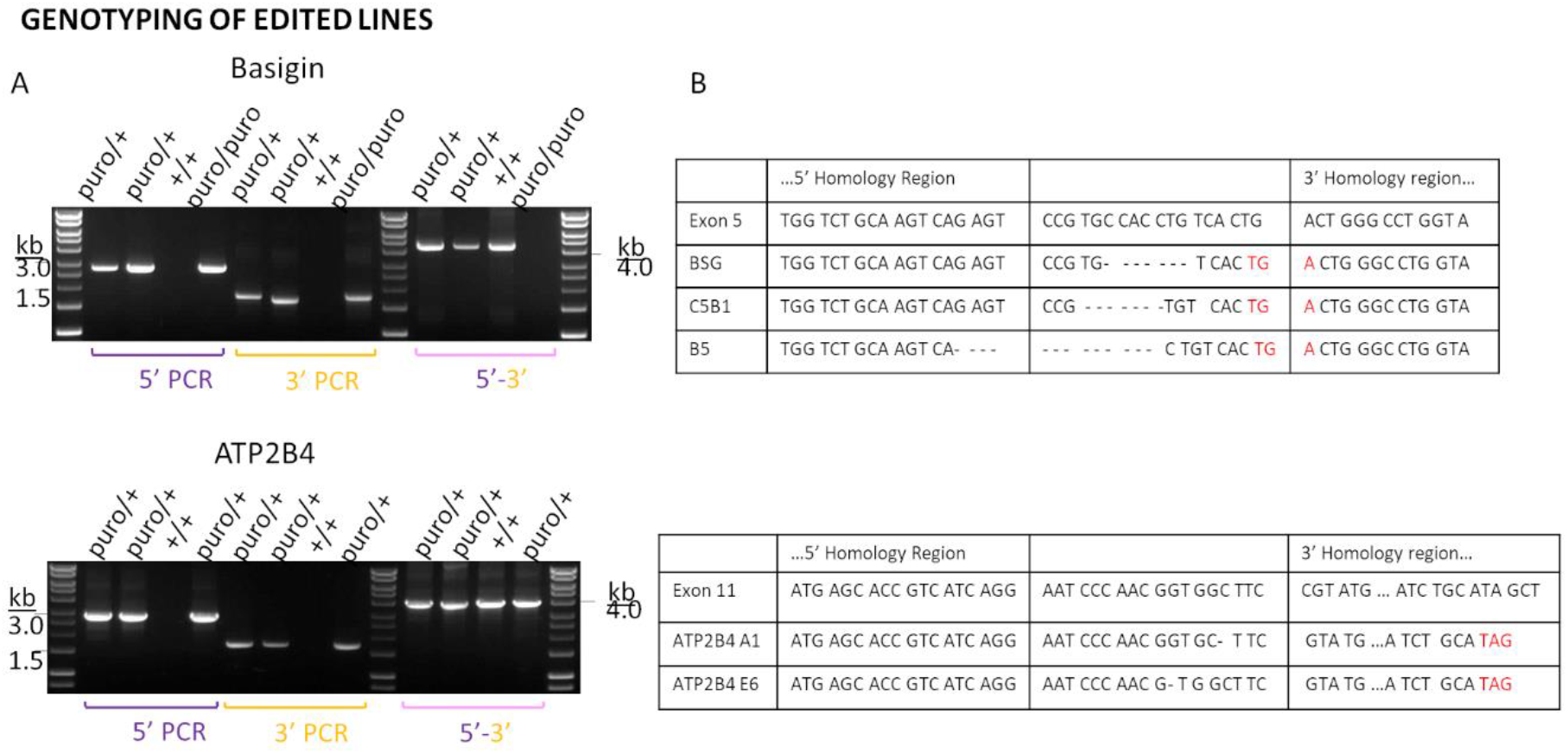
Verification of edited cell lines. The seams of the homology arms’ boundaries were tested by PCR for insertion of the cassette. Sequencing through the second allele verified disruption and confirmed deletion.

## Author Contributions

**Alena Pance, Bee Ling, Kioko Mwikali, Manousos Koutsourakis, Chukwuma Agu, Foad Rouhani, Hannes Ponstingl, Ruddy Montandon, Frances Law, Julian C. Rayner**

AP conception and design of the project, performing of experiments and writing of the manuscript; JCR conception of the project and writing of the manuscript; BL setting up and running of flow cytometry approach. KM bioinformatics analysis; HP handling of the data and analysis; MK genome editing of BSG and ATP2B4; CA and FR repogramming of iPSC lines; RM help and guidance with flow cytometric analysis; FL help and advice for stem cell culture and maintenance.

## REFERENCES

1. C. Aurrecoechea et al., PlasmoDB: a functional genomic database for malaria parasites. Nucleic Acids Res 37, D539–543 (2009).

2. T. W. Lo et al., Precise and heritable genome editing in evolutionarily diverse nematodes using TALENs and CRISPR/Cas9 to engineer insertions and deletions. Genetics 195, 331–348 (2013).

3. F. A. Ran et al., Genome engineering using the CRISPR-Cas9 system. Nat Protoc 8, 2281–2308 (2013).

4. M. Ghorbal et al., Genome editing in the human malaria parasite Plasmodium falciparum using the CRISPR-Cas9 system. Nat Biotechnol 32, 819–821 (2014).

5. J. Straimer et al., Site-specific genome editing in Plasmodium falciparum using engineered zincfinger nucleases. Nat Methods 9, 993–998 (2012).

6. A. F. Cowman, C. J. Tonkin, W. H. Tham, M. T. Duraisingh, The Molecular Basis of Erythrocyte Invasion by Malaria Parasites. Cell Host Microbe 22, 232–245 (2017).

7. G. E. Weiss et al., Revealing the sequence and resulting cellular morphology of receptor-ligand interactions during Plasmodium falciparum invasion of erythrocytes. PLoS Pathog 11, e1004670 (2015).

8. A. F. Cowman, D. Berry, J. Baum, The cellular and molecular basis for malaria parasite invasion of the human red blood cell. J Cell Biol 198, 961–971 (2012).

9. D. Damena, A. Denis, L. Golassa, E. R. Chimusa, Genome-wide association studies of severe P. falciparum malaria susceptibility: progress, pitfalls and prospects. BMC Med Genomics 12, 120 (2019).

10. N. Malaria Genomic Epidemiology, Insights into malaria susceptibility using genome-wide data on 17,000 individuals from Africa, Asia and Oceania. Nat Commun 10, 5732 (2019).

11. S. N. Kariuki, T. N. Williams, Human genetics and malaria resistance. Hum Genet 139, 801–811 (2020).

12. A. K. Bei, C. Brugnara, M. T. Duraisingh, In vitro genetic analysis of an erythrocyte determinant of malaria infection. J Infect Dis 202, 1722–1727 (2010).

13. C. Crosnier et al., Basigin is a receptor essential for erythrocyte invasion by Plasmodium falciparum. Nature 480, 534–537 (2011).

14. E. S. Egan et al., Malaria. A forward genetic screen identifies erythrocyte CD55 as essential for Plasmodium falciparum invasion. Science 348, 711–714 (2015).

15. K. Trakarnsanga et al., An immortalized adult human erythroid line facilitates sustainable and scalable generation of functional red cells. Nat Commun 8, 14750 (2017).

16. T. J. Satchwell et al., Genetic manipulation of cell line derived reticulocytes enables dissection of host malaria invasion requirements. Nat Commun 10, 3806 (2019).

17. E. J. Scully et al., Generation of an immortalized erythroid progenitor cell line from peripheral blood: A model system for the functional analysis of Plasmodium spp. invasion. Am J Hematol, (2019).

18. I. International Stem Cell et al., Characterization of human embryonic stem cell lines by the International Stem Cell Initiative. Nat Biotechnol 25, 803–816 (2007).

19. A. Veres et al., Low incidence of off-target mutations in individual CRISPR-Cas9 and TALEN targeted human stem cell clones detected by whole-genome sequencing. Cell Stem Cell 15, 27–30 (2014).

20. D. Hendriks, H. Clevers, B. Artegiani, CRISPR-Cas Tools and Their Application in Genetic Engineering of Human Stem Cells and Organoids. Cell Stem Cell 27, 705–731 (2020).

21. K. Takahashi et al., Induction of pluripotent stem cells from adult human fibroblasts by defined factors. Cell 131, 861–872 (2007).

22. F. A. Soares, R. A. Pedersen, L. Vallier, Generation of Human Induced Pluripotent Stem Cells from Peripheral Blood Mononuclear Cells Using Sendai Virus. Methods Mol Biol 1357, 23–31 (2016).

23. C. A. Agu et al., Successful Generation of Human Induced Pluripotent Stem Cell Lines from Blood Samples Held at Room Temperature for up to 48 hr. Stem Cell Reports 5, 660–671 (2015).

24. F. Rouhani et al., Genetic background drives transcriptional variation in human induced pluripotent stem cells. PLoS Genet 10, e1004432 (2014).

25. H. Kilpinen et al., Common genetic variation drives molecular heterogeneity in human iPSCs. Nature 546, 370–375 (2017).

26. M. Carrasquilla et al., Defining multiplicity of vector uptake in transfected Plasmodium parasites. Sci Rep 10, 10894 (2020).

27. S. H. Adjalley, M. C. Lee, D. A. Fidock, A method for rapid genetic integration into Plasmodium falciparum utilizing mycobacteriophage Bxb1 integrase. Methods Mol Biol 634, 87–100 (2010).

28. R. Frita et al., Simple flow cytometric detection of haemozoin containing leukocytes and erythrocytes for research on diagnosis, immunology and drug sensitivity testing. Malar J 10, 74 (2011).

29. A. T. Y. Yeung et al., Exploiting induced pluripotent stem cell-derived macrophages to unravel host factors influencing Chlamydia trachomatis pathogenesis. Nat Commun 8, 15013 (2017).

30. M. E. Ritchie et al., limma powers differential expression analyses for RNA-sequencing and microarray studies. Nucleic Acids Res 43, e47 (2015).

31. L. Vallier et al., Early cell fate decisions of human embryonic stem cells and mouse epiblast stem cells are controlled by the same signalling pathways. PLoS One 4, e6082 (2009).

32. M. Rebelo et al., A novel flow cytometric hemozoin detection assay for real-time sensitivity testing of Plasmodium falciparum. PLoS One 8, e61606 (2013).

33. C. Timmann et al., Genome-wide association study indicates two novel resistance loci for severe malaria. Nature 489, 443–446 (2012).

34. S. J. Lu et al., Biologic properties and enucleation of red blood cells from human embryonic stem cells. Blood 112, 4475–4484 (2008).

35. S. Hirose et al., Immortalization of erythroblasts by c-MYC and BCL-XL enables large-scale erythrocyte production from human pluripotent stem cells. Stem Cell Reports 1, 499–508 (2013).

36. T. J. Egan, Haemozoin formation. Mol Biochem Parasitol 157, 127–136 (2008).

37. A. J. Chen et al., Quantitative imaging of intraerythrocytic hemozoin by transient absorption microscopy. J Biomed Opt 25, 1–11 (2019).

38. C. Delahunt, M. P. Horning, B. K. Wilson, J. L. Proctor, M. C. Hegg, Limitations of haemozoin-based diagnosis of Plasmodium falciparum using dark-field microscopy. Malar J 13, 147 (2014).

39. C. M. Ndila et al., Human candidate gene polymorphisms and risk of severe malaria in children in Kilifi, Kenya: a case-control association study. Lancet Haematol 5, e333–e345 (2018).

40. S. Lessard et al., An erythroid-specific ATP2B4 enhancer mediates red blood cell hydration and malaria susceptibility. J Clin Invest 127, 3065–3074 (2017).

41. M. G. Dalghi et al., Plasma membrane calcium ATPase activity is regulated by actin oligomers through direct interaction. J Biol Chem 288, 23380–23393 (2013).

42. X. Gao, K. Gunalan, S. S. Yap, P. R. Preiser, Triggers of key calcium signals during erythrocyte invasion by Plasmodium falciparum. Nat Commun 4, 2862 (2013).

43. J. C. Volz et al., Essential Role of the PfRh5/PfRipr/CyRPA Complex during Plasmodium falciparum Invasion of Erythrocytes. Cell Host Microbe 20, 60–71 (2016).

44. M. L. Gazarini, A. P. Thomas, T. Pozzan, C. R. Garcia, Calcium signaling in a low calcium environment: how the intracellular malaria parasite solves the problem. J Cell Biol 161, 103–110 (2003).

45. B. Zambo et al., Decreased calcium pump expression in human erythrocytes is connected to a minor haplotype in the ATP2B4 gene. Cell Calcium 65, 73–79 (2017).

46. S. S. Ho et al., Microsatellite markers within --SEA breakpoints for prenatal diagnosis of HbBarts hydrops fetalis. Clin Chem 53, 173–179 (2007).

47. R. Hong, U. Chandola, L. F. Zhang, Cat-D: a targeted sequencing method for the simultaneous detection of small DNA mutations and large DNA deletions with flexible boundaries. Sci Rep 7, 15701 (2017).

48. S. M. Taylor, C. Cerami, R. M. Fairhurst, Hemoglobinopathies: slicing the Gordian knot of Plasmodium falciparum malaria pathogenesis. PLoS Pathog 9, e1003327 (2013).

49. V. Pathak, R. Colah, K. Ghosh, Effect of inherited red cell defects on growth of Plasmodium falciparum: An in vitro study. Indian J Med Res 147, 102–109 (2018).

50. R. Huisjes et al., Squeezing for Life - Properties of Red Blood Cell Deformability. Front Physiol 9, 656 (2018).

51. A. Bunyaratvej, P. Butthep, N. Sae-Ung, S. Fucharoen, Y. Yuthavong, Reduced deformability of thalassemic erythrocytes and erythrocytes with abnormal hemoglobins and relation with susceptibility to Plasmodium falciparum invasion. Blood 79, 2460–2463 (1992).

52. S. Glushakova et al., Hemoglobinopathic erythrocytes affect the intraerythrocytic multiplication of Plasmodium falciparum in vitro. J Infect Dis 210, 1100–1109 (2014).

